# Evolving properties of biological materials captured via needle-based cavity expansion method

**DOI:** 10.1101/2024.09.23.614613

**Authors:** Hannah M. Varner, S. Kiana Naghibzadeh, Katie C. Spaeth, Abigail Klein, Tal Cohen

## Abstract

**Background:** The mechanical properties of biological tissues change over time and with disease progression. Quantifying these mechanical properties can thus be instrumental for medical diagnosis and for evaluation of tissue viability for transplant. However, soft and biological materials are exceptionally challenging to mechanically characterize using conventional testing methods, which are hindered by limitations of sample size, fixturing capabilities, and sample preparation.

**Objective:** We hypothesize that Volume Controlled Cavity Expansion (VCCE) is well-suited to capture subtle mechanical differences in biological tissue. The objective of this work is therefore twofold: first, we seek to quantify how stiffness of liver and gelatin evolve with age. In achieving this understanding, we aim to demonstrate the precision of VCCE in measuring subtle changes in the mechanical properties of biological tissues.

**Methods:** Performing VCCE tests over 15 days in samples of gelatin and liver (porcine and bovine), we track the evolving pressure-volume response and deformation limits of the materials.

**Results:** In both materials, we observed time-dependent variation of the stiffness and fracture thresholds. In gelatin VCCE repeatably captured stiffening over time, which was correlated with a higher fracture stress. This was in contrast to observations in bovine liver, where stiffening corresponded to a lower fracture stress. Porcine liver initially stiffened, then reversed this trend and relaxed.

**Conclusion:** Through this work we show that liver and gelatin stiffen with age, and that this trend is measurable via VCCE. These results highlight the utility of VCCE and call attention to the need for a new class of mechanism based constitutive models that are capable of capturing variations in material over time with a minimal number of parameters.

## 1. Introduction

The mechanical properties of biological tissues change over time and in different states of health and disease. A precise mechanical testing method has the potential to quantify evolution of these properties and inform both diagnoses and predictive models.

Connecting mechanical properties to clinically relevant information is an enduring practice, with manual palpation going back millennia to doctors in ancient Egypt [1], and lasting into modern medical practice as scientists attempt to quantify these results [2]. This is with good reason: a soft blood clot may either dissolve, or calcify and stiffen over time [3]; during rapid tumor growth the densification of cells and changes to the extracellular matrix increase stiffness enough that it can be felt via manual palpation [4]; as liver fibrosis progresses, connective tissue accumulates in the extracellular matrix causing stiffening [5, 6, 7, 8, 9, 10]; malignant thyroid nodules may be stiffer than benign nodules [11]; arterial stiffness is a predictor of morbidity in renal disease [12, 13]; and a decrease in stiffness of a transplant kidney within the first month is the typical progression [14].

Mechanical properties of organs are also crucial in understanding injury and survivability. In motor vehicle crash testing, whole organs from human cadavers or large mammals are used to evaluate damage and inform protective measures [15, 16]. But just as with meat brought home from a supermarket, how these organs are stored prior to testing may change their mechanical properties. Hence, the outcomes of crash testing may not be fully representative of an in vivo response of a human experiencing a traumatic injury. Multiple authors have noted that evidence is inconclusive on how freezing and refrigeration affect the stiffness of ex vivo organs, and how this varies by species or tissue [10, 17, 18].

Despite a clear need for, and interest in, cataloging these properties and correlating in vivo and ex vivo data [19, 20], challenges arise in both the process of measuring and quantifying tissue properties. With the development of new testing methodologies, mechanical testing has the potential to be a minimally invasive diagnostic method that would not require any expensive laboratory assays outside of the clinic, and could rapidly evaluate changes undergone by tissue in storage.

The ability to perform several targeted tests within an organ in its native environment is required to build accurate models. Nanoindendation testing is nondestructive, however it cannot reach internal target tissues in vivo [21], meanwhile, conventional compression, tension or rheological testing require modifications (shaping, cutting into a tensile dog bone or a compression cube) to perform tests on bulk samples. Targeting and slice selection in non-invasive methods like Magnetic Resonance Imaging (MRI) and Elastography (MRE) are subjective [9] or require inference about the nature of complex heterogeneity to interpret stiffness of an internal feature [22]. Further, though in use as a diagnosis tool in liver fibrosis [23], MRE is limited by its mathematical assumptions to be predictive of only small, linear, tissue deformations, when nonlinear behavior is characteristic of most biological materials. Additionally, outside of the body, the boundary conditions of a sample may interfere with the accuracy of a measurement, and the changes in stiffness may be more subtle and difficult to register.

Proposed alternative methods that aim to overcome these challenges such as needle induced cavitation rheology (NICR) [24, 25, 26] are also unable to determine a nonlinear response. Laser induced cavitation combined with high speed imaging has been compared with predictions of nonlinear elastic and viscoelastic models for identification of material properties [27], but requires preparation and visualization conditions that restrict it to in-vitro testing [28]. An alternative method proven in ex vivo tissue is Volume Controlled Cavity Expansion (VCCE), which has been previously used to measure properties of both synthetic [29, 30, 31] and biological [32, 25] materials. Material stretch is controlled through the volume of an expanding cavity, thus allowing interpretation of nonlinear material behavior, and continued observation throughout stable fracture propagation and relaxation. VCCE can be deployed in-situ internal to a material sample and provides measurement localized to the millimeter scale.

A common way to quantify material stiffness and predict how these materials will respond under load is using a continuum framework with nonlinear constitutive models such as those developed by Ogden [33], Mooney-Rivlin [34, 35], Yeoh [36], and Fung [2]. Acknowledging the inhomogeneity of biological tissues, many efforts have been made to develop constitutive models that capture additional complexity [37, 38, 39, 40, 41, 42, 43], however, the amount of data required to fully inform these models [44] has not resulted in widespread use. Even within the limited subset of “simple” constitutive models, there is not consensus as to which model is best suited for various biological materials. This is highlighted by several authors in their evaluation of various internal organs [19, 45, 46, 47], and by the authors of this work for whole blood clots [32].

*With the primary goal of demonstrating how VCCE, as a minimally invasive method, can be used to measure even subtle mechanical changes of tissue over time*, in this paper, we measure local properties in the bulk of biological tissues as they age^1^. We use gelatin as a baseline biologically-derived material, then progress to measuring porcine and bovine liver to assess results in cell-based materials. All of these materials are valuable to understand in their own right, and this provides the basis of our secondary objective: to evaluate the changes in each that occur over time.

Gelatin is the most abundant animal protein [48] and has thus been used as a tissue mimic by many authors [49, 21, 50, 51, 52, 53, 54]. Liver is uniquely susceptible to damage during trauma and motor vehicle accidents because of its size and location [55]. Bovine and porcine liver serve as common surrogates to human liver in crash tests [56, 57], and the need to store organs between tests has motivated authors to evaluate how stiffness changes after removing the organ from the body [5], and after being stored either refrigerated or frozen [15, 17, 18].

This paper is organized as follows: Section 2 describes the tissue types and handling, experimental procedure, and analytical frameworks used. In Section 3, we detail the results of aging studies in both gelatin and liver, describe how conventional constitutive models are not well-suited to model the phenomena we observe, then discuss how these results are influenced by sample weight. We finish the analysis by examining the critical values indicative of a transition between elastic expansion and damage in Section 3.5 and concluding remarks in Section 4.

## 2. Methods

Two model systems of contrasting complexity are used in this work: gelatin and liver, as discussed in Section 2.1.

These are tested using VCCE, discussed in Section 2.2, with the data analysis described in Section 2.3.

### 2.1. Materials used and their structure

Gelatin is a chemically modified form of collagen, the main component of animal connective tissue, and can be made to have elastic properties that are reported to be similar to porcine and bovine liver and other organs [51, 52, 58, 59, 60]. Gelatin bonds are thermally reversible and organized in a combination of amorphous, disordered coils, and ordered regions of triple helices with a typical length scale on the order of 15 nm [48, 58]. In this study, commercial 225 bloom porcine gelatin was prepared at a volume water *v* to weight dry gelatin *w* ratio of *v* : *w* ratio of 10:1 (9% *w/v*). Gelatin powder was bloomed with half cold distilled water, then dissolved and stirred for 5 minutes with the remaining half of the distilled water heated to boiling. Samples from the same batch were molded into multiple containers and refrigerated between testing. All replicates *n* reported for a given test day are from a single batch, across multiple containers.

Liver, by contrast, contains cells that are organized into a defined tissue structure. The typical cell size within bovine liver is 20 µm with ducts up to ∼1 millimeter, which are organized into lobules on the order of 2-3 mm as illustrated in Fig. 1(a). Lobules in porcine liver are segmented by collagenous septa, while lobules in bovine liver do not contain significant collagen boundaries and are more similar to the structure found in human liver [56]. Bovine liver was obtained frozen from a commercial butcher. Porcine liver was collected from adult female Yorkshire swine then sectioned into individual samples (∼ 3×3×3 cm) and refrigerated (0-4 °C) within 30 minutes postmortem.

**Figure 1:**
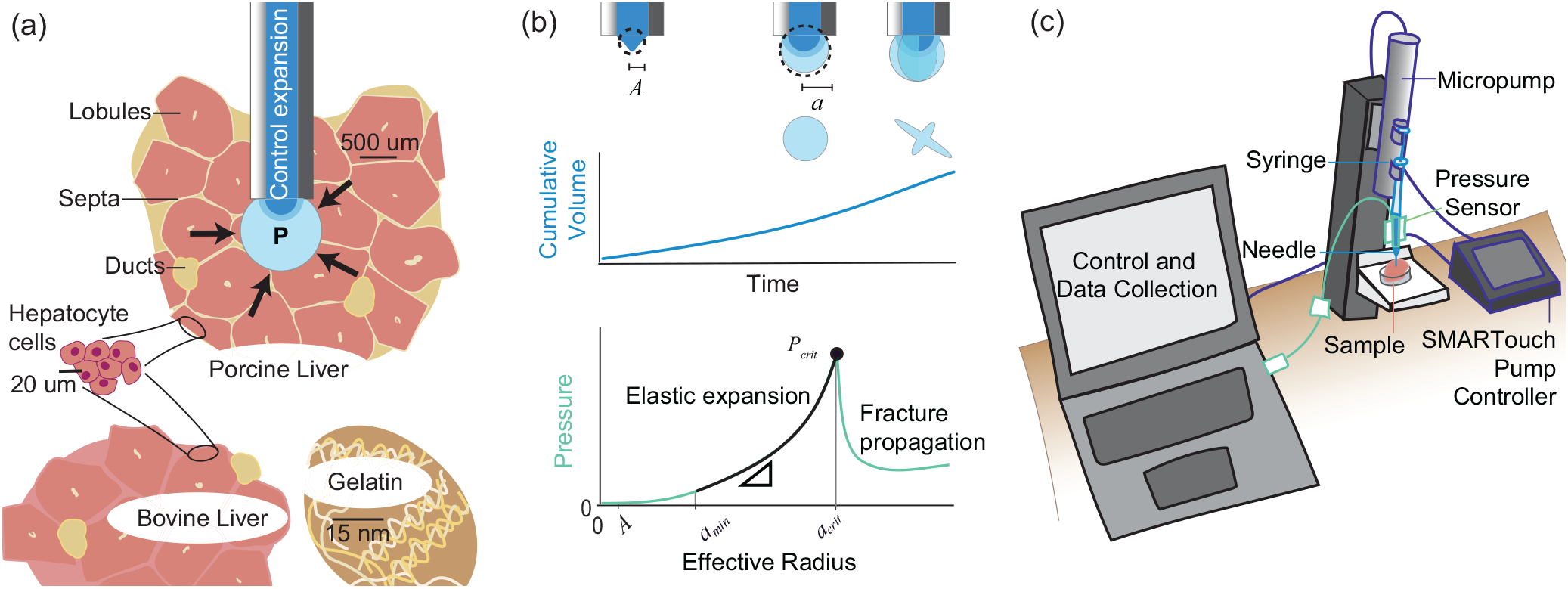
(a) The sample is penetrated by a blunt needle and the tissue structure is assumed to be well approximated by a continuum. Controlling the expansion of a cavity over time, (b) VCCE returns a pressure due to material resistance to deformation that is characterized by an initial elastic region, followed by a critical pressure *P*_*crit*_ and equivalent radius *a*_*crit*_. (c) This data is collected via a custom software script that dictates the expansion profile programmed into the SMARTouch controller.

The effect of tissue storage protocol on stiffness was one of the variables investigated in this work (as discussed in Sec. 3.4). Therefore, tissues were stored in either an air tight plastic bag or submerged in phosphate buffered saline (PBS) depending on the protocol of a given trial as delineated in Table 1. For each day of testing, a single sample was removed from refrigeration and allowed to warm to room temperature prior to testing (with *n* replicates performed within the same sample). All experiments using animal-derived tissue were approved by and performed in accordance with the University Committee on Animal Care.

**Table 1:** Sample descriptions and identifiers. All samples were refrigerated between tests.

| Species | Tissue | Origin | Storage | Abbreviation | $n$ |
| --- | --- | --- | --- | --- | --- |
| Bovine | Liver | Commercial butcher. Received frozen, defrosted. | Airtight bag | Liver B1 | 42 |
|  |  |  | PBS | Liver B2 | 25 |
| Porcine | Liver | Adult, female, Yorkshire swine research animal. Collected fresh then refrigerated. | *Split*: PBS and air-tight bag | Liver P1 | 48 |
|  |  |  | PBS | Liver P2 | 25 |
| Porcine | Gelatin | Commercial gelatin. Fabricated in house, refrigerated. | Closed container | Gel P1 | 21 |
|  |  |  | Closed container | Gel P2 | 24 |

### 2.2. Cavity initiation and control

In a VCCE test, a needle is inserted into the target region of a tissue. After allowing for material relaxation, a millimeter-scale cavity is expanded using incompressible fluid, as previously described [32, 29, 31, 61]. To interpret the response of the material, the pressure *P* is measured and correlated to the effective radius *a* (calculated assuming the cavity is spherical) throughout the controlled expansion. Fig. 1(b) shows a schematic of the typical response whereby the pressure increases until a critical value *P*_*crit*_ is reached at the corresponding radius *a*_*crit*_. The critical values are assumed to be indicative of the onset of damage in the material [62].

For this work, VCCE is performed at a constant volume rate (CVR) expansion^2^ of 600 nL/s using a World Precision Instruments (WPI) UltraMicroPump3 controlled via a custom script input to a WPI SMARTouch Controller as detailed by Unikewicz et al. [61] and shown schematically in Fig. 1(c). The MicroPump is fitted with a 10 µL Hamilton syringe body, enabling a maximum cavity size of *a* =1.3 mm. A 25 gauge, 1 inch, blunt, luer lock needle is inserted 12-15 mm then retracted 2-4 mm to allow for relaxation of the sample to *P <* 5 kPa. Cavity expansion follows immediately after the relaxation. Pressure is recorded via a PendoTech PRESS-S-000 flow-through pressure sensor. PBS is used as the working fluid in order to maintain isotonic balance with the tissues.

### 2.3. Data interpretation and analysis

To avoid imposing bias from the damage process after *P*_*crit*_ on the interpretation of the elastic expansion, all equation fitting takes place from *a*_*min*_ = 0.153 mm to *a*_*crit*_ (Fig. 1(b)) ^3^. We assume that all tests on a given day with a material return results from the same population of mechanical data and perform a nonlinear least squares fit in MATLAB with each test weighted equally (an example is provided in Appendix Appendix A). The root mean square error (RMSE) is calculated to evaluate the quality of all fits.

Four expressions were selected to relate *P* and *a*, three empirical expressions and one physics-based constitutive model. As the data are observed to be strain stiffening, the equations chosen all have an increasing first derivative to capture the curvature of the data. The three empirical models are: an exponential *P* = *c*_1_ exp(*c*_2_*a*) + *c*_3_, a polynomial *P* = *c*_1_*a*^2^ + *c*_2_*a* + *c*_3_, and a power function 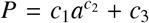. The final fitting option employs a 1-term Ogden model, which can be rewritten in spherical coordinates with the strain energy density of *W* = 2*µ/α*^2^(2*λ*^*α*^ + *λ*^−2*α*^ −3) and integrated as in [29] to return a relation between pressure and volume. The three fit parameters in the Ogden model are the shear modulus *µ*, a stiffening parameter *α*, and an initial defect size of effective radius *A* (shown on Fig. 1(b)) that determines the stretch *λ* = *a/A*.

## 3. Results

Performing VCCE in the materials of interest over a number of days (Fig. 2) we see that the method is able to repeatably measure subtle changes that indicate the effect of storage method and time on the mechanical properties of gelatin and liver.

**Figure 2:**
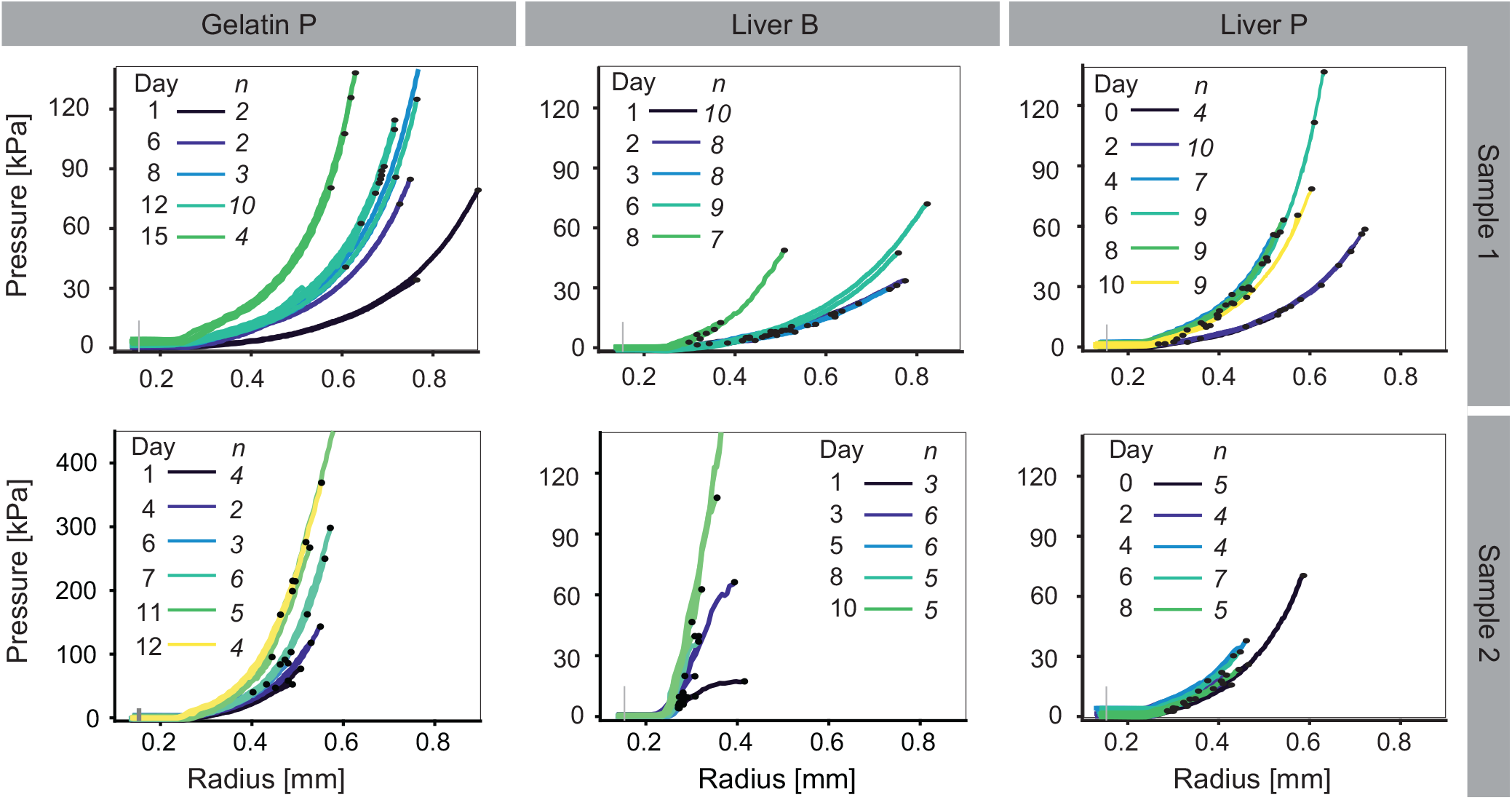
Raw data for *n* tests performed in the same samples each day demonstrate qualitative stiffening with age in all samples. Critical values of individual tests are marked with a • and the grey vertical line indicates the lower bound of data that are used for fitting. The first column presents two samples of porcine gelatin tested over 15 days (Gel P1) and 12 days (Gel P2), the second column presents previously frozen bovine liver over 8 or 10 days (Liver B1 and B2) and the rightmost column presents fresh porcine liver tested over 10 days (Liver P1) and 8 days (Liver P2).

### 3.1. Stiffening measurements are repeatable

To quantify how the mechanical properties evolve, we first examine the raw data returned from the VCCE measurements. Fig. 2 captures the pressure-radius data collected for gelatin, bovine liver and porcine liver samples (from left to right, respectively). Within the same test day, multiple observations *n* are seen to overlap well, indicating precision in the measurements. A majority of the tests show clear strain stiffening (i.e. a non-linear *P*-*a* relationship), and all samples demonstrate change in stiffness with age.

### 3.2. Conventional constitutive models are insufficient

In order to quantify the observed stiffening and track changes over time, our goal is to describe the data using a model. Traditionally, this would be done with a constitutive model relating deformations to loads via material parameters. However, as previously mentioned, there is minimal agreement as to the optimal constitutive model to use for both bovine and porcine tissue. Previous works have chosen to use a range of models including: 1-to 3-term Ogden [15, 17, 63], 1-to 2-term Mooney-Rivlin [47], low- and high-strain stiffness [47], linear [6, 18], neo-Hookean [23], Fung [20], Yeoh [63], custom metrics [5], or a combination of established model terms [42]. Additionally, a number of researchers have highlighted the difficulty of using the Ogden model fit under one set of loading conditions to predict response in other loading scenarios [64, 65]. Therefore, we also investigated fitting several purely empirical models (polynomial, exponential, and power function). All models, empirical or physics based, have three parameters and are fit as described in Sec. 2.3 with the RMSE used to evaluate “best fit”.

Empiric fits, not a conventional constitutive model, produced lower error and better convergence for a majority of tests. Table A.2 of Appendix Appendix A details of fit parameters by day for all samples. In gelatin, the two samples are consistently fit by different models (power function for Gel P1 and exponential for Gel P2). In liver, the best fit for the experimental data varies by sample and age between a polynomial and power function.

Overall, our data are poorly fit (if at all) by both the 1-term Ogden model, and fitting with physically unrealistic parameters occurs in half of the test days. Manual tuning of the bounds, start points and fitting region of the nonlinear least squares fitting algorithm was required to produce Ogden material parameter sets that converge within the domain. A consistent set of these inputs could not be found that would result in a successful fit across all days of testing for the 1-term Ogden model applied to gelatin or liver. Based on previously reported shear moduli obtained using an Ogden model for porcine gelatin [59, 66, 67], bovine liver [17, 15], and porcine liver [60, 20], *µ <* 0.100 kPa are deemed unrealistic. An initial defect size of *A <* 0.025 mm is similarly considered unrealistic given this is less than 1/10^th^ the radius of the needle inserted in the sample and previous work by the authors found *A* in whole blood clots was typically on the order of 0.10 mm [32]. Additional discussion of the fitting bounds can be found in Appendix Appendix A.

The current models are insufficient to concisely capture the complexity of changes occurring in biological tissue.

In order to fit the evolving mechanical response of the tissues observed in our results, we anticipate that relying on empiric equation as a foundation will likely require fewer overall terms than conventional constitutive models.

### 3.3. Stiffness increases with storage, to a point

In the absence of a conventional constitutive model, we propose using the tangent to the curve in order to understand how material stiffness changes with age. Examining the raw data in Fig. 2, the relationship of *P* and *a* is non-linear and therefore a single number cannot be used to describe the stiffness at all deformations. We choose to define a “tangent stiffness” *k* as the first derivative of the curve, multiplied by the inner diameter of the needle (0.26 mm) with resulting units of kPa. To compare the stiffness of a sample as it ages, we examine *k* at the same pressure for subsequent days. We compute *k* of the best fitting (lowest RMSE) model relating *P* and *a* at ten regular pressure intervals between 5 and 105 kPa and plot this over time in Fig. 3.

**Figure 3:**
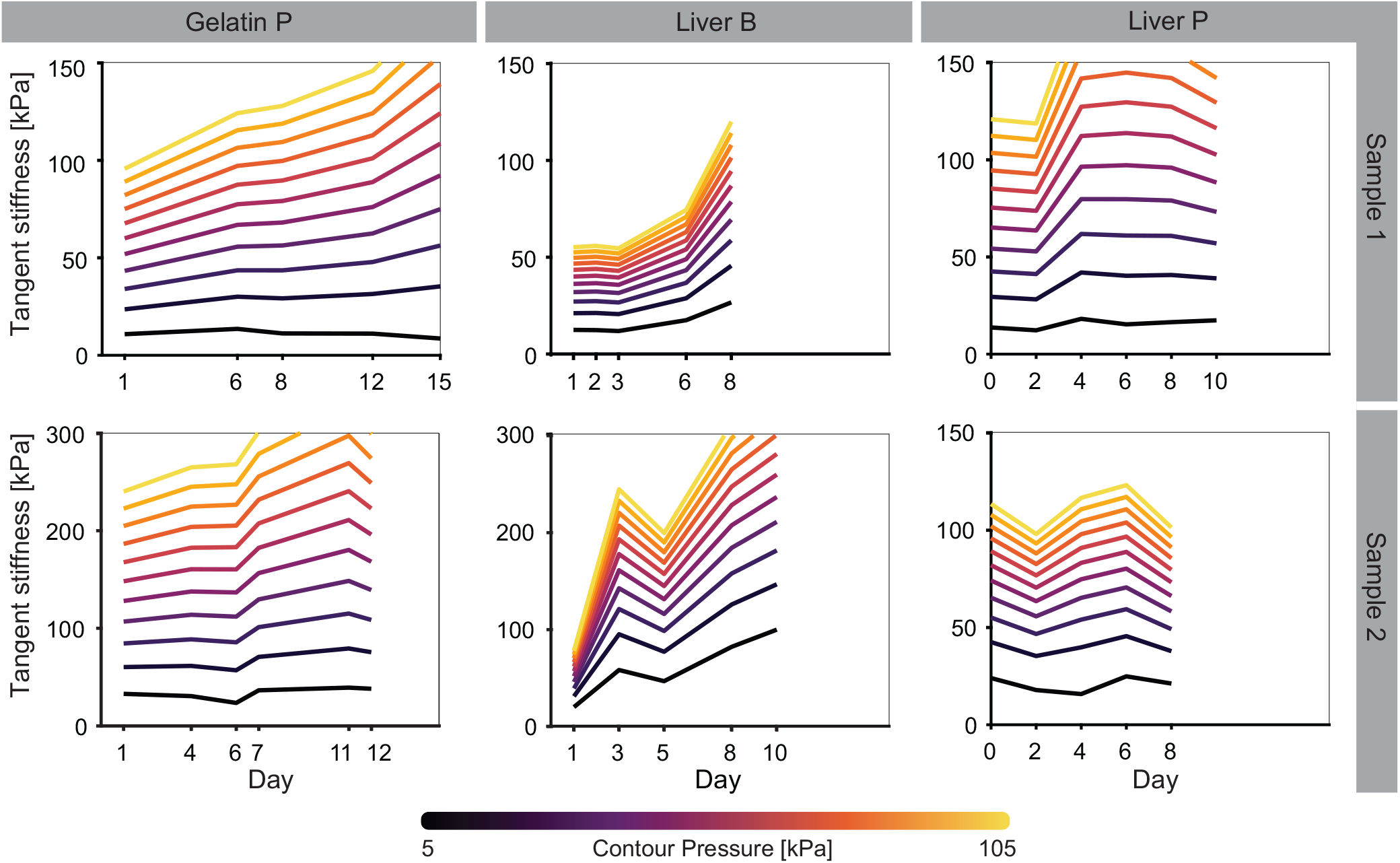
Tangent stiffness *k* for a fixed pressure taken every 10 kPa starting at 5 kPa and plotted relative to test day shows increased stiffening in gelatin and bovine liver and a stiffening followed by softening in porcine liver. Slight non-monotonic stiffening is observed in the second sample of gelatin and bovine liver. Stiffness are calculated from the model form with the lowest RMSE across all days for each material.

Tangent stiffness increases through Day 8 in a majority of both liver and gelatin samples. In all samples, the changes are less dramatic at low pressures, corresponding to the small strain (linear) regime. The highest overall tangent stiffness values are observed in the second gelatin and bovine liver samples. Bovine liver stiffens more rapidly than gelatin or porcine liver, displaying the steepest contour lines across all pressures.

A number of samples display non-monotonic changes in pressure with age. This occurred most notably across both porcine liver samples, where *k* decreases in the final day of testing. Previous work by Lu et al. [18] observed stiffening of porcine liver samples refrigerated for 20 days, however samples were not tracked over time, so it is not possible to know the evolution within the test window.

### 3.4. Stiffness increase cannot be explained by mass loss in all samples

While tissues were stored in air-tight bags or submerged in PBS, gelatin samples were in sealed containers with one surface exposed to air, potentially allowing for variable degrees of water loss between samples. If the water content of samples varied over time, the resulting material properties may have also changed. At lower water content the collagen helices in gelatin may rearrange into more ordered, stiffer structures. Likewise, a higher solid mass fraction in dehydrated liver tissue could lead to densification and increasing stiffness. By weighing samples each day, we aim to understand whether mass loss is connected to the stiffness changes observed in Fig. 3.

Fig. 4 shows the ratio of final to initial mass *m/m*_0_ for select gelatin and liver samples. Dehydration of the gelatin is apparent, with mass losses of 0.16% per day in Gel P1 and 0.31% per day in Gel P2. By contrast, consistent mass loss is not observed in the liver samples, and yet we still record stiffening for these materials. While not ruled out in gelatin, mass loss cannot be the sole mechanism causing stiffening over time in our samples.

**Figure 4:**
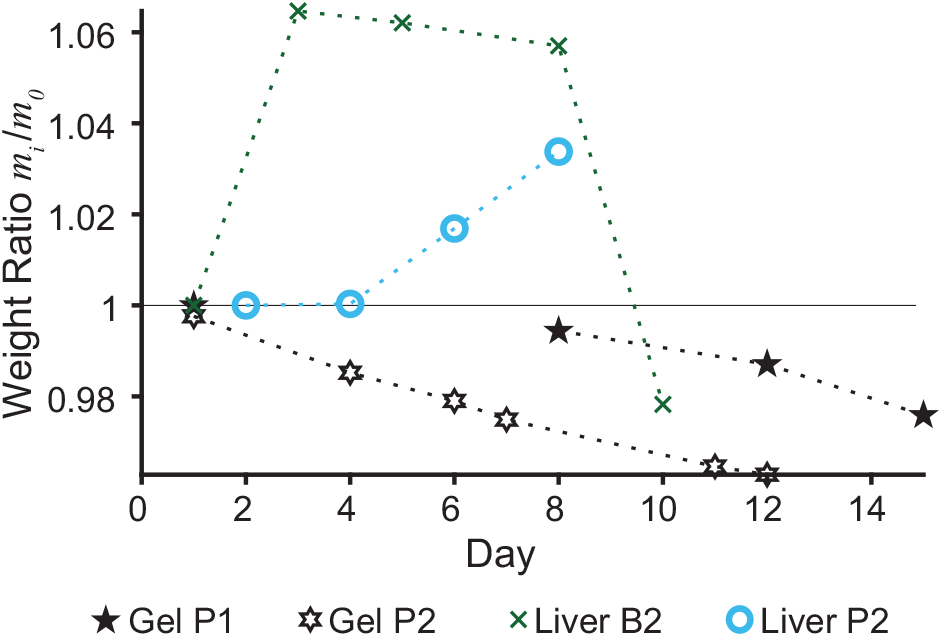
The ratio of initial to final sample weight *m*_*i*_*/m*_0_. Tracking this in gelatin and liver over time shows that weight loss is inversely proportional to the stiffness increase observed in Fig. 3 for gelatin (black stars), however the quantities are not tightly connected in porcine liver (blue circles) and bovine liver (green crosses).

Investigating storage method further, no significant differences were noted between the stiffness measurements of porcine liver samples from the same animal stored in airtight bags compared to those submerged in PBS. Fig. 5 shows multiple pressure-radius curves of two samples obtained from the same animal and tested in parallel over the course of 10 days. Results are repeatable for each test day across multiple trials, regardless of the storage method used.

**Figure 5:**
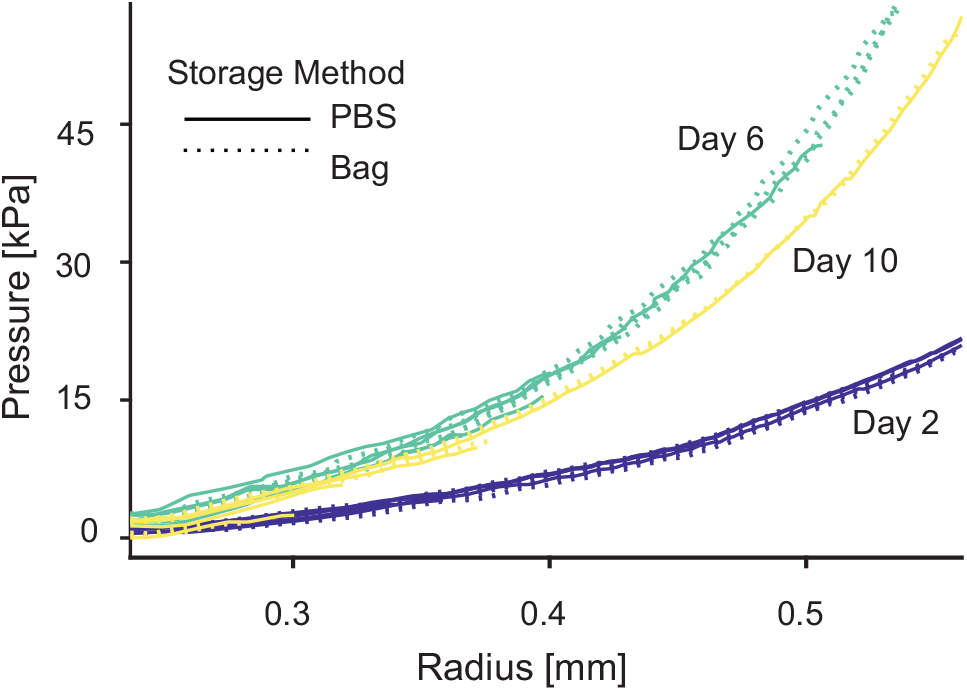
Comparing the pressure-volume signatures of porcine liver samples stored in PBS and sealed bags shows indistinguishable pressure-radius curves between the two storage methods.

### 3.5. Decreasing trend in a_crit_ over time is not matched in P_crit_ for all materials

The critical values are indicative of the damage threshold in the material over time. All materials except Liver B2 show a decrease in the median critical radius 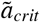 over time (Fig. 6). In all cases showing decreasing *a*_*crit*_, the value declined by over 25% by the end of testing. Results with a statistically significant difference between days (at *p<* 0.05) are noted alongside their *p* value.

**Figure 6:**
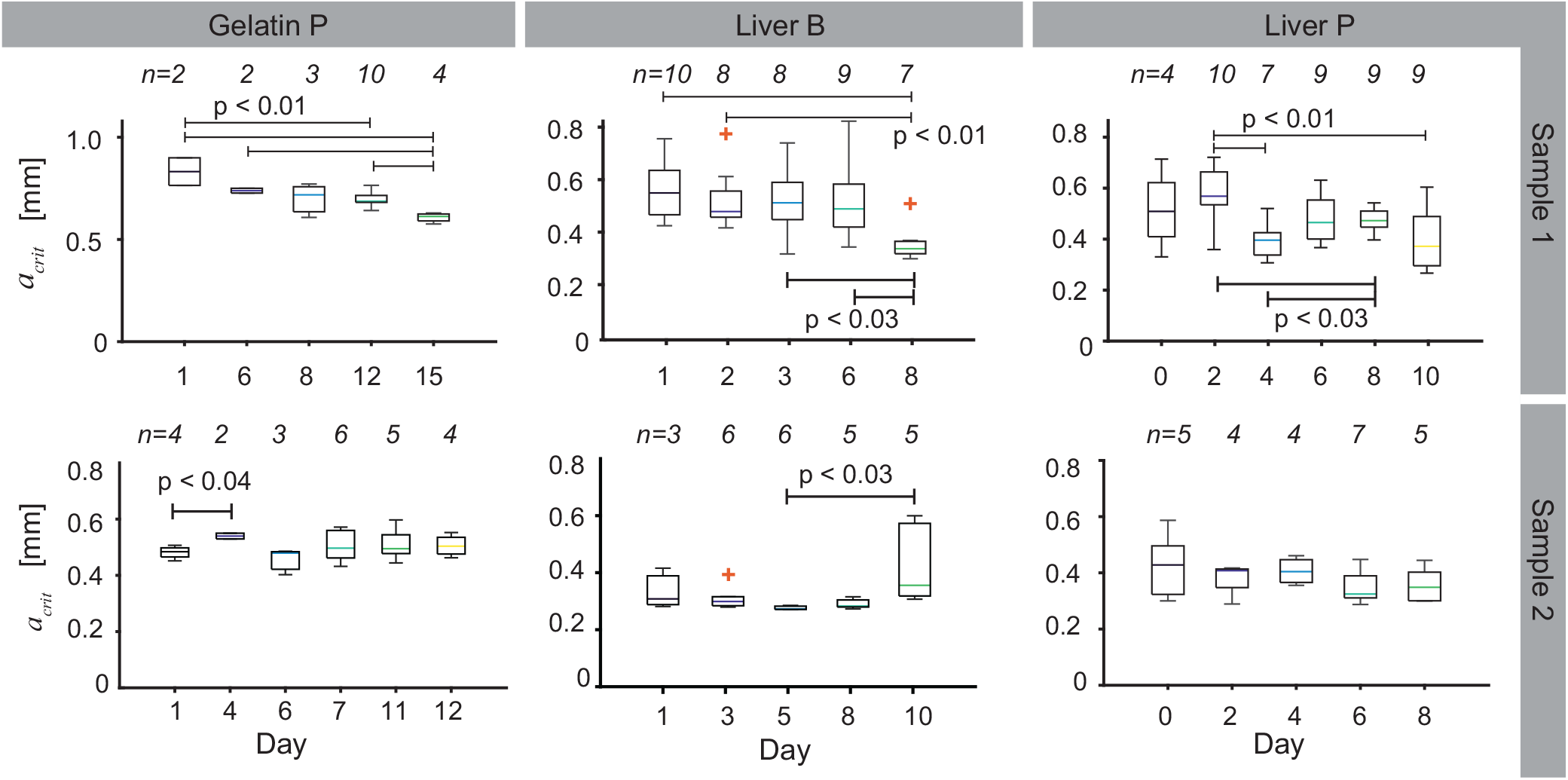
Critical radius for gelatin, bovine liver and porcine liver displayed as box plots. Box extends between the first and third quartile (*q*_1_ and *q*_3_) with the colored center line at the median. Whiskers extend to the furthest data point that is *<* 1.5(*q*_3_ − *q*_1_) and outliers beyond this are marked with red crosses. Number of samples (*n*) within each day are listed above each chart and samples with statistically significant distributions are indicated along with the *p* value.

Gelatin demonstrated a monotonically increasing median critical pressure 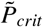 that more than doubled after 15 and 12 days of storage (Fig. 7 Sample Gel P1 and Gel P2). By contrast, 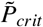 in bovine and porcine liver do not increase monotonically or clearly over the course of 8-10 days (Fig. 7 Liver B1, B2 and P1, P2). The reversal of elastic behavior at day 10 observed for porcine liver in Fig. 3 is not obvious within the critical value trends, emphasizing that different mechanisms govern elasticity of a material verses the fracture and damage.

**Figure 7:**
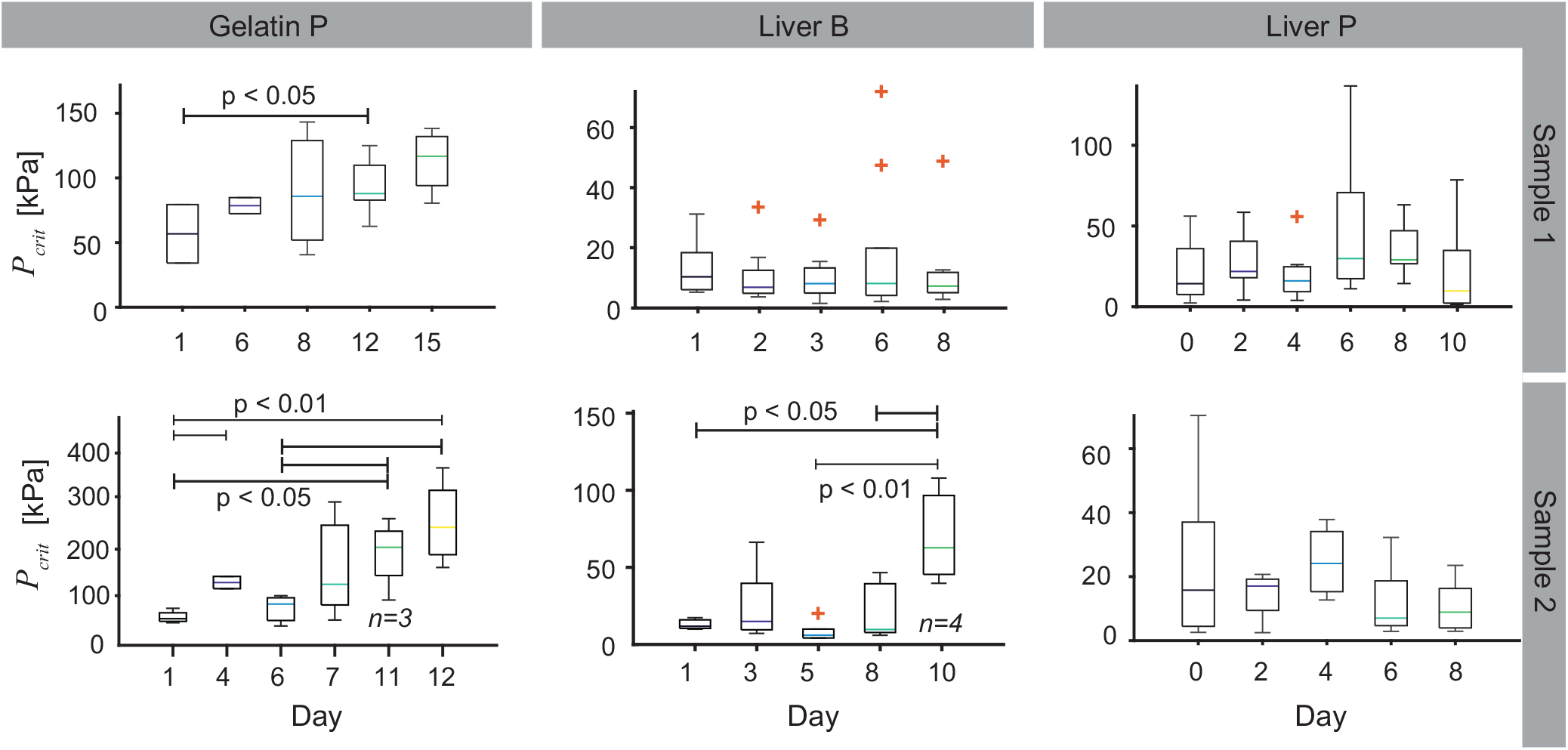
Critical pressure for gelatin increases with age (Gelatin P1 and P2), while there is no trend with age in bovine liver (Liver B1 and B2) and porcine liver (Liver P1 and P2). Box extends between the first and third quartile (*q*_1_ and *q*_3_) with the colored center line at the median. Whiskers extend to the furthest data point that is *<* 1.5(*q*_3_ − *q*_1_) and outliers beyond this are marked with red crosses. Number of samples (*n*) are the same as those listed in Fig. 6 except where specified, indicating a measurement outside of the calibration range of the pressure sensor. Samples with statistically significant distributions are indicated along with the *p* value.

The freshly collected porcine liver samples tested here had a generally higher *P*_*crit*_ than the previously frozen bovine samples with comparable *a*_*crit*_. In their tension testing of dog bone shaped liver samples, Kemper et al. [56] saw that failure strains of porcine liver are higher than those of human liver, attributing the difference to thicker collegenated septa between the lobules that are present in porcine liver. Defined septa are absent from bovine liver as well and may explain the discrepancy seen in our data. Ultimate strain in liver is commonly found to be between 25% and 35% [47, 68, 69, 70, 71, 72] with ultimate stresses between 1.85 and 10.9 kPa [69, 71]. Lu et al. [17] observed in testing bovine liver, that the failure stress increased while the failure strain decreased for samples frozen for 30 or 60 days. This aligns with the lower *a*_*crit*_ observed in bovine liver in our results as well, however an increase in *P*_*crit*_ as a surrogate measure of ultimate strength is not as clear.

## 4. Conclusions

In this work, we have shown that VCCE is fully capable of capturing subtle biological changes over time. As VCCE has been scaled down to a bench top device [61], this has opened up the possibility for its use in the clinical setting as a powerful indicator of tissue normalcy and health. Additional studies with larger sample sizes will be required in order to achieve the goals of understanding organ viability for transplant, drawing comparisons between the responses of fresh versus stored organs in destructive testing, or drawing population-level conclusions about diagnosis. However, with VCCE removing the barrier to sensitive, local, in vivo measurements, these are no longer the question in collecting sufficient data, simply the availability of resources to do so.

Many hyperelastic constitutive models were first developed for synthetic materials and remain valuable in that domain. However, when applied to biological materials there remains room for improvement. Here we show how important this is in the context of tissue aging as the empiric expression that provide the best fit to our data evolved on subsequent days. There is a risk of over fitting these models by blindly applying additional terms in order to capture this temporal aging. We propose that empirical expressions are sufficient to capture changes in behavior. Nonetheless, future work should consider the mechanistic changes in tissue structure that take place during aging. A new class of models is required to describe the time dependent biological changes with minimal fitting parameters through thoughtfully incorporating terms that capture the specific timescales of degradation within the inhomogeneous cellular, molecular and fibrous components that make up complex tissue hierarchies.

Just as Reiter et al. [73] turned to the microstructure of brain matter as a way to explain viscoelasticity observed therein, this could inform time dependent aging as well. Likewise, Seyedpour et al. [45] discuss the characteristic length scales of tissue microstructure and pores as integral factors in understanding and measuring fibrotic liver tissue via MRI. After death, autolysis of cells occurs within the first 36 hours [74, 75]. This process is driven by enzymatic activity native to cells, which breaks down proteins and ruptures cell walls. Liver is a metabolically active organ, leading to high concentrations of enzymes [19, 76] and relatively faster rates of autolysis than other organs in the body [77]. While refrigeration delays these processes, the damage to cell wall and tissue protein may explain the non-monotonic change in tangent stiffness observed here in porcine liver. By contrast, freezing denatures different enzymes at different rates [78]. The breakdown of enzymes in the frozen bovine liver, resulting in reduced enzyme activity after defrosting, may also explain why the same response was not observed in the bovine samples. Tissue integrity postmortem and the resulting measured stiffness is clearly linked to the biological changes that occur within it, however more research is required to directly understand how individual components of the tissues contribute to stiffness, and to link the varying rates at which these decay to mechanical responses.

While the experimental technique described in this work was applied to a limited subset of biological tissues, VCCE is versatile enough to be applied across a range of other material systems. Its ability to measure local mechanical properties at the millimeter scale in soft materials overcomes previous challenges with sample geometry and fixturing that impede progress in testing many soft materials.

## Funding and Acknowledgements

The authors thank Prof. Giovanni Traverso, Niora Fabian DVM, Andrew Pettinari and the DCM for support in sample collection. We acknowledge the partial support of our work through Office of Naval Research grant N000142312530. H.V. acknowledges the support of the Department of Defense (DOD) through the National Defense Science & Engineering Graduate (NDSEG) Fellowship Program. S.K.N. acknowledges the MIT Postdoctoral Fellowship Program for Engineering Excellence (PFPFEE).

## Authorship

All authors contributed to the study conception and design. Material preparation, and data collection were performed by Hannah Varner, S. Kiana Naghibzadeh, Katie Spaeth and Abigail Klein. Analysis was performed by Hannah Varner. The first draft of the manuscript was written by Hannah Varner and all authors commented on previous versions of the manuscript. All authors read and approved the final manuscript.

## Appendix A. Fitting

Fitting is performed by regularizing all the tests performed on that day with an equal number of sample points between *a*_*min*_=0.153 mm and *a*_*crit*_ then using a non-linear least squares algorithm to fit the model equation. The lower bound at *a*_*min*_ is informed via the same geometric assumptions described previously by the authors in [32]. Given the variability of *a*_*crit*_, this may result in the fit being biased to low radii, however this is justified as there is more rich information in that region. Fig. A.8 shows an example of four fit types on Day 15 for sample Gel P1.

**Figure A.8:**
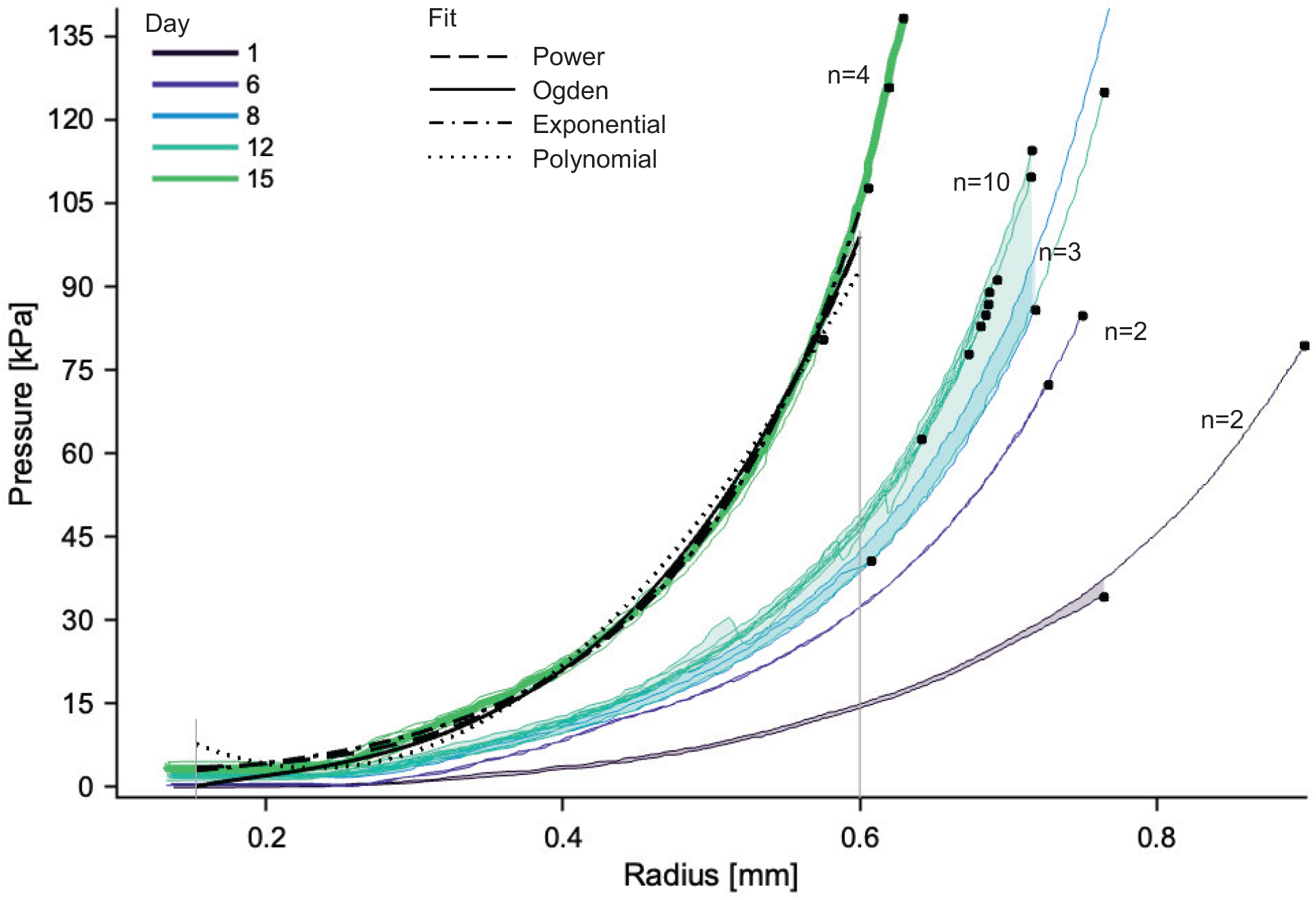
VCCE data collected in gelatin (Gel P1) over 15 days. Each line is a test performed on a given day with the shaded area between curves capturing the spread. Critical values are marked with a black circle. Representative fitting on Day 15 is the result of the nonlinear least squares fit for the four listed models.

Table A.2 includes the RMSE value for the fits across all days in gelatin, bovine liver and porcine liver. The lowest RMSE where fitting converged to physically realistic parameters is highlighted in bold for each day. Note that while the Ogden model at times produced a lower RMSE fit than other models, the resulting fitting parameters were physically unrealistic in these cases and therefore not selected as the ideal fit for the experimental data. The lower bound of *A* was 0.025 mm, as this is 1/10^th^ the diameter of the needle and our previous work in whole blood clots [32] would indicate that the initial damage created via needle insertion is greater than this limit. Typical *µ* reported in literature range from 11.34-29.9 kPa in porcine gelatin [59, 66, 67], 12.7-39.7 kPa in bovine liver [17, 15], 0.30 kPa-1.63 MPa in porcine liver [60, 20]. Therefore, fit *µ* that returned the lower bound of the fitting domain (0.100 kPa) are at least an order of magnitude less than a majority of the previously published data and determined to be unrealistic as well. Sample Liver B2 did not converge well with the domain for Days 3, 5, and 8 using an exponential or polynomial model as well.

**Table A.2:** Fitting parameters for expressions as applied to gelatin, bovine liver and porcine liver

| | $P = c_1 e^{c_2 a} + c_3$ | | | | $P = c_1 a^{c_2} + c_3$ | | | | $P = c_1 a^2 + c_2 a + c_3$ | | | | $\Psi = 2\mu/\alpha^2(2\lambda^\alpha + \lambda^{-2\alpha} - 3)$ | | | |
| --- | --- | --- | --- | --- | --- | --- | --- | --- | --- | --- | --- | --- | --- | --- | --- | --- |
| | $c_1$ | $c_2$ | $c_3$ | RMSE | $c_1$ | $c_2$ | $c_3$ | RMSE | $c_1$ | $c_2$ | $c_3$ | RMSE | $\mu$ | $A=a/\lambda$ | $\alpha$ | RMSE |
| Gel P1 |  |  |  |  |  |  |  |  |  |  |  |  |  |  |  |  |
| Day 1 | 0.648 | 5.37 | -2.06 | **0.425** | 93.3 | 3.62 | -0.167 | 0.436 | 192 | -118 | 17.8 | 2.55 | 0.575 | 0.182 | 7.34 | 0.811 |
| Day 6 | 1.16 | 5.74 | -3.57 | **0.823** | 223 | 3.71 | -0.054 | 1.03 | 298 | -152.6 | 19.8 | 2.29 | 1.24 | 0.169 | 7.09 | 1.10 |
| Day 8 | 0.729 | 6.80 | -0.93 | **1.14** | 261 | 3.70 | 1.32 | 1.21 | 451 | -254 | 36.9 | 5.12 | 1.28 | 0.174 | 7.48 | 2.96 |
| Day 12 | 0.896 | 6.68 | -1.58 | **0.997** | 355 | 3.98 | 1.69 | 1.09 | 419 | -219 | 30.1 | 3.60 | 1.37 | 0.155 | 7.04 | 2.64 |
| Day 15 | 0.923 | 7.89 | -0.67 | **1.78** | 807 | 4.17 | 3.16 | 2.07 | 725 | -350 | 45.3 | 4.43 | 2.45 | 0.158 | 7.28 | 2.75 |
| Gel P2 |  |  |  |  |  |  |  |  |  |  |  |  |  |  |  |  |
| Day 1 | 0.570 | 9.94 | -6.77 | 2.53 | 1850 | 4.59 | -3.51 | **2.37** | 868 | -378 | 38.2 | 2.52 | 2.26 | 0.181 | 8.89 | 3.26 |
| Day 4 | 0.696 | 9.81 | -7.01 | 1.65 | 2740 | 4.92 | -2.09 | **1.59** | 1340 | -635.6 | 72.3 | 4.42 | 2.42 | 0.173 | 8.65 | 2.18 |
| Day 6 | 0.511 | 11.0 | -3.38 | 3.145 | 3220 | 4.84 | 0.126 | **3.03** | 1250 | -548 | 60.5 | 3.39 | 0.100 | 0.075 | 7.89 | 2.84 |
| Day 7 | 0.580 | 10.9 | -7.67 | 1.91 | 4730 | 5.17 | -2.85 | **1.70** | 2280 | -1130 | 134 | 9.80 | 2.63 | 0.175 | 9.19 | 3.19 |
| Day 11 | 0.584 | 12.1 | -8.07 | 5.27 | 1.04E4 | 5.45 | -2.28 | **3.80** | 3490 | -1710 | 202 | 13.05 | 0.100 | 0.080 | 8.54 | 6.27 |
| Day 12 | 1.29 | 10.5 | -11.6 | 3.02 | 7190 | 4.94 | -2.33 | **2.49** | 3260 | -1530 | 175 | 9.72 | 0.100 | 0.069 | 8.09 | 2.90 |
| Liver B1 |  |  |  |  |  |  |  |  |  |  |  |  |  |  |  |  |
| Day 1 | 0.801 | 5.23 | -2.91 | 0.634 | 83.9 | 3.17 | -1.05 | 0.544 | 107 | -45.3 | 4.43 | **0.454** | 0.100 | 0.094 | 6.68 | 0.770 |
| Day 2 | 0.422 | 6.31 | -1.77 | 0.477 | 84.5 | 3.23 | -0.763 | 0.428 | 110 | -47.5 | 5.01 | **0.322** | 0.100 | 0.092 | 6.61 | 0.62 |
| Day 3 | 0.356 | 6.36 | -1.33 | 0.449 | 90.1 | 3.53 | -0.323 | **0.359** | 105 | -46.9 | 5.29 | 0.512 | 0.100 | 0.097 | 6.72 | 0.549 |
| Day 6 | 0.591 | 5.95 | -2.79 | 0.664 | 158.2 | 3.89 | -0.984 | **0.545** | 193.6 | -98.3 | 11.64 | 1.33 | 0.641 | 0.180 | 7.52 | 1.00 |
| Day 8 | 0.261 | 10.6 | -2.09 | 0.895 | 1076 | 4.50 | -0.653 | 0.760 | 506 | -210 | 21.6 | **0.734** | 0.100 | 0.090 | 7.93 | 0.87 |
| Liver B2 |  |  |  |  |  |  |  |  |  |  |  |  |  |  |  |  |
| Day 1 | 4.64 | 4.71 | -10.9 | 2.04 | 225 | 2.48 | -3.15 | **1.93** | 211 | -32.5 | -0.85 | 1.98 | 3.19 | 0.16 | 6.94 | 2.28 |
| Day 3 | 0.001 | 31.9 | -0.10 | 1.09 | 1.12E6 | 9.37 | 0.153 | 0.924 | 2090 | -819 | 79.2 | **3.29** | 0.100 | 0.079 | 8.79 | 4.33 |
| Day 5 | 0.001 | 23.7 | 0.10 | 0.444 | 5.00E9 | 16.4 | 0.139 | 0.392 | 1390 | -552 | 53.8 | **1.93** | 0.100 | 0.176 | 23.1 | 1.44 |
| Day 8 | 0.001 | 31.7 | -0.36 | 1.12 | 1.91E9 | 15.1 | 0.161 | 0.716 | 3360 | -1340 | 132 | **4.31** | 0.100 | 0.127 | 13.9 | 4.12 |
| Day 10 | 10.5 | 7.73 | -52.2 | 13.2 | 5220 | 3.54 | -20.3 | 10.9 | 4250 | -1590 | 145 | **6.63** | 0.235 | 0.050 | 7.13 | 15.15 |
| Liver P1 |  |  |  |  |  |  |  |  |  |  |  |  |  |  |  |  |
| Day 0 | 0.771 | 6.08 | -2.85 | 0.549 | 191 | 3.76 | -0.443 | **0.516** | 213 | -99.6 | 11.6 | 1.06 | 0.100 | 0.091 | 7.00 | 0.584 |
| Day 2 | 0.755 | 6.11 | -2.13 | 0.553 | 185 | 3.74 | 0.274 | **0.498** | 208 | -98.4 | 12.3 | 1.04 | 0.540 | 0.136 | 6.91 | 0.511 |
| Day 4 | 0.665 | 8.71 | -2.60 | 1.38 | 804 | 4.06 | 0.109 | 1.26 | 499 | -201 | 21.4 | **1.24** | 0.100 | 0.068 | 7.09 | 1.20 |
| Day 6 | 0.623 | 8.53 | -1.80 | 0.933 | 828 | 4.29 | 1.05 | **0.820** | 682 | -325 | 39.8 | 3.10 | 1.45 | 0.152 | 7.66 | 1.26 |
| Day 8 | 0.648 | 8.58 | -2.14 | 1.37 | 793 | 4.16 | 0.650 | 1.36 | 493 | -207 | 23.1 | 1.66 | 0.941 | 0.128 | 7.29 | **1.34** |
| Day 10 | 0.818 | 7.65 | -3.19 | 1.10 | 566 | 3.99 | -0.165 | **1.01** | 441 | -188 | 20.5 | 1.31 | 0.100 | 0.075 | 7.14 | 0.951 |
| Liver P2 |  |  |  |  |  |  |  |  |  |  |  |  |  |  |  |  |
| Day 0 | 0.711 | 7.95 | -3.17 | 1.37 | 630 | 4.13 | -0.343 | **1.24** | 455 | -198 | 21.8 | 1.50 | 0.100 | 0.078 | 7.26 | 1.21 |
| Day 2 | 0.628 | 8.63 | -1.71 | 2.13 | 479 | 3.60 | 0.325 | 2.08 | 344 | -124 | 12.8 | **1.84** | 0.124 | 0.060 | 6.55 | 1.91 |
| Day 4 | 0.448 | 9.84 | 0.15 | 1.05 | 925 | 4.12 | 2.15 | 0.963 | 494 | -194 | 22.2 | **0.830** | 1.128 | 0.109 | 6.67 | 1.15 |
| Day 6 | 0.109 | 13.4 | -0.67 | 1.33 | 1575 | 4.70 | 0.039 | 1.28 | 539 | -220 | 23.1 | **1.14** | 0.100 | 0.081 | 7.72 | 1.28 |
| Day 8 | 0.378 | 9.66 | -1.98 | 1.08 | 647 | 4.00 | -0.425 | 1.02 | 364 | -139 | 13.65 | **0.844** | 0.100 | 0.080 | 7.44 | 0.978 |

## Footnotes

1 Note, we use the term “aging” not in the sense of *biological age* since the start of growth, but rather time since collection or acquisition of a tissue sample. This may either be by tissue harvest from a specimen animal, or from the point of entry into the laboratory (defrosting in the case of commercially obtained samples, or fabrication of samples made in the lab).

2 Previous work with VCCE has highlighted the benefit of performing constant radial expansion rate, and therefore a constant stretch rate, in order to determine viscoelastic mechanical constants tied to constitutive models [31, 32]. CVR ensures a rate saturation early in the expansion profile of the cavity, therefore reliably accessing the instantaneous shear modulus *µ*_*inst*_ even of viscoelastic samples such as liver [17].

3 The lower bound is determined per the same rationale used in [32] and the upper bound is determined to exclude higher values that may bias the fit given the nonlinearity of the data.

